# A Neanderthal Origin of Genetic Variants Associated with Asthma

**DOI:** 10.1101/2023.10.30.564624

**Authors:** Andrew S. H. Day, Kyle J. Gaulton

## Abstract

While genome-wide association studies have implicated a number asthma-associated genes, whether the disease-linked variants originate from ancient genomes has not been explored. Here we show that of the 51 asthma-associated loci that we surveyed, 39 carry variants that were derived in the Neanderthal lineage. The shared sequences suggest that some asthma variants may have originated from the Neanderthal genome after admixture and subsequent introgression into the Eurasian population. Of note, one variant, rs4742170, previously linked to asthma and childhood wheezing, was shown in a recent study to disrupt glucocorticoid receptor binding to a putative *IL33* enhancer, and elevate enhancer activity of this key asthma gene. Further analysis of the *IL33*-associated variant in publicly available single nucleus ATACseq data of the human lung addressed its localization in open chromatin, providing an additional approach to screen for variants that may impact the expression of asthma-associated genes. Together, these findings suggest a possible Neanderthal origin of genetic variants associated with asthma.

## INTRODUCTION

Neanderthals migrated out of Africa and inhabited Eurasia before their extinction. Their interbreeding with anatomically modern humans has left a lasting genetic legacy in our genomes. The admixture dating back to approximately 60,000 years ago was evident through comparison of Neanderthal genomes with modern human genomes (1). Furthermore, Neanderthal-derived single nucleotide polymorphisms (SNPs) are found at higher frequencies in current Eurasian populations compared to Yoruba African populations (2), consistent with an admixture event occurring outside of Africa. It is estimated that non-African modern humans share approximately 1-4% of their DNA with Neanderthals (1, 3). Among these variants, many can contribute to traits including those related to immunity, metabolism, and external features (4, 5).

One of the most notable areas where Neanderthal genes have had a significant impact is the human immune system. It was postulated that Neanderthals had already adapted to the pathogens present in Eurasia, and interbreeding allowed modern humans to inherit beneficial genetic variants for immune defense (6, 7). This conferred advantages to modern humans as they encountered new pathogens during their migration and thereafter (8, 9). Of note, recent studies reveal that both a COVID-19 genetic risk factor and a genetic protective factor were introduced to humans from Neanderthals (10, 11). On the other hand, the same introduction of Neanderthal sequences also brought variants associated with autoimmune diseases. Neanderthal-derived single nucleotide polymorphisms (SNPs) have been associated with Crohn’s disease and lupus (12-16). These autoimmune diseases reflect the trade-off between immunity against pathogens and the potential for immune overactivity.

Like these autoimmune diseases, asthma is associated with heightened immune response. Whether genetic risk of asthma has an origin in Neanderthals has been challenging to discern due to the complex causes of asthma. Recent genome-wide association studies (GWAS) have identified genetic variants linked to asthma susceptibility in modern humans (17-20). Whether these variants have their origin in Neanderthals has not been extensively evaluated. In this study, we found that a subset of asthma-associated loci carry variants that are present in the Neanderthal genome, but not Yoruba African populations, providing evidence that they were inherited during admixture events after Neanderthal migration out of Africa.

## RESULTS

### Evidence for asthma variants in the Neanderthal genome

We used a curated list of 51 non-HLA loci that have been associated with asthma risk (21). To elevate the strength of the evidence for an origin in Neanderthals, we adopted a previously-used approach, most recently to demonstrate the link between COVID-19 genetic risk and Neanderthals (2, 4, 11). Using this approach in our study, we analyzed genomic regions carrying asthma risk variants in three specific populations, the Vindija Neanderthal ancient genome, the French genome (HGDP00533) and the Yoruba genome (HGDP00936) from the 1000 Genomes Project. The Vindija genome was selected because it is postulated to be the Neanderthal genome most like the population that had the initial contact with modern humans migrating out of Africa (22, 23). The Yoruba genome was selected because there was no admixture between Yoruba and Neanderthal populations after Neanderthal migration out of Africa, as supported by the little to no post-migration Neanderthal-derived allele in the Yoruba genome (2, 23). The French genome was selected to represent Eurasian modern human populations post migration out of Africa. We searched for genetic signatures that are present in both the Vindija and French genomes but absent in the Yoruba genome, as an indicator that these variants were inherited from Neanderthals as a result of Neanderthal-human admixture.

Among the 51 loci analyzed, 39 of them were found to contain variants shared between the Vindija and French genomes, but are not present in the Yoruba genome (Table 1). We also observed that the Neanderthal derived alleles in the French genome tend to cluster together (Figure 1). This is likely due to linkage disequilibrium, and represents the Neanderthal haplotypes that exist in the Eurasian population (24-26). Together, the shared sequences suggest that these asthma variants in modern humans were derived from the Neanderthal lineage because of introgression after interbreeding.

**Table 1.**
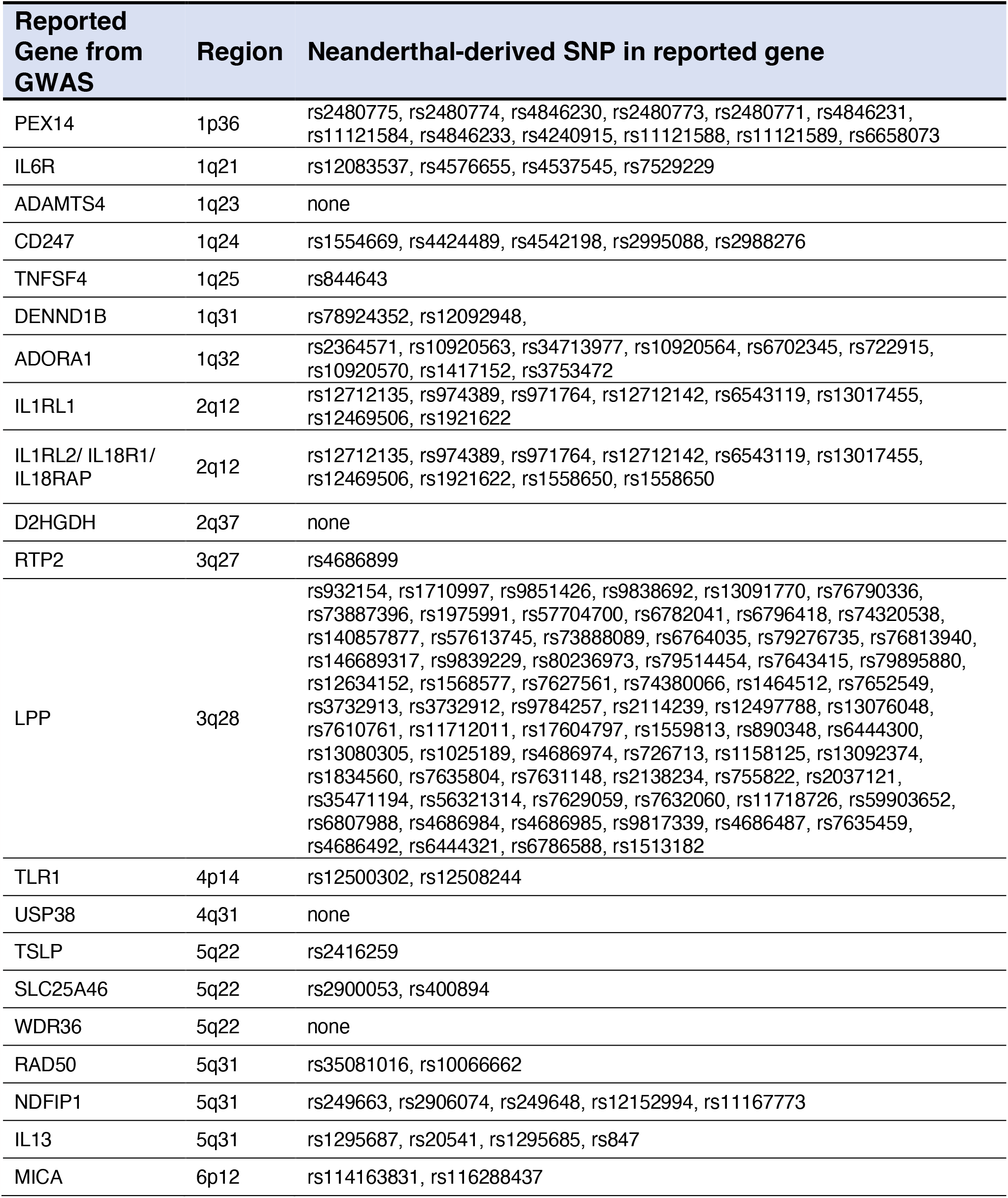
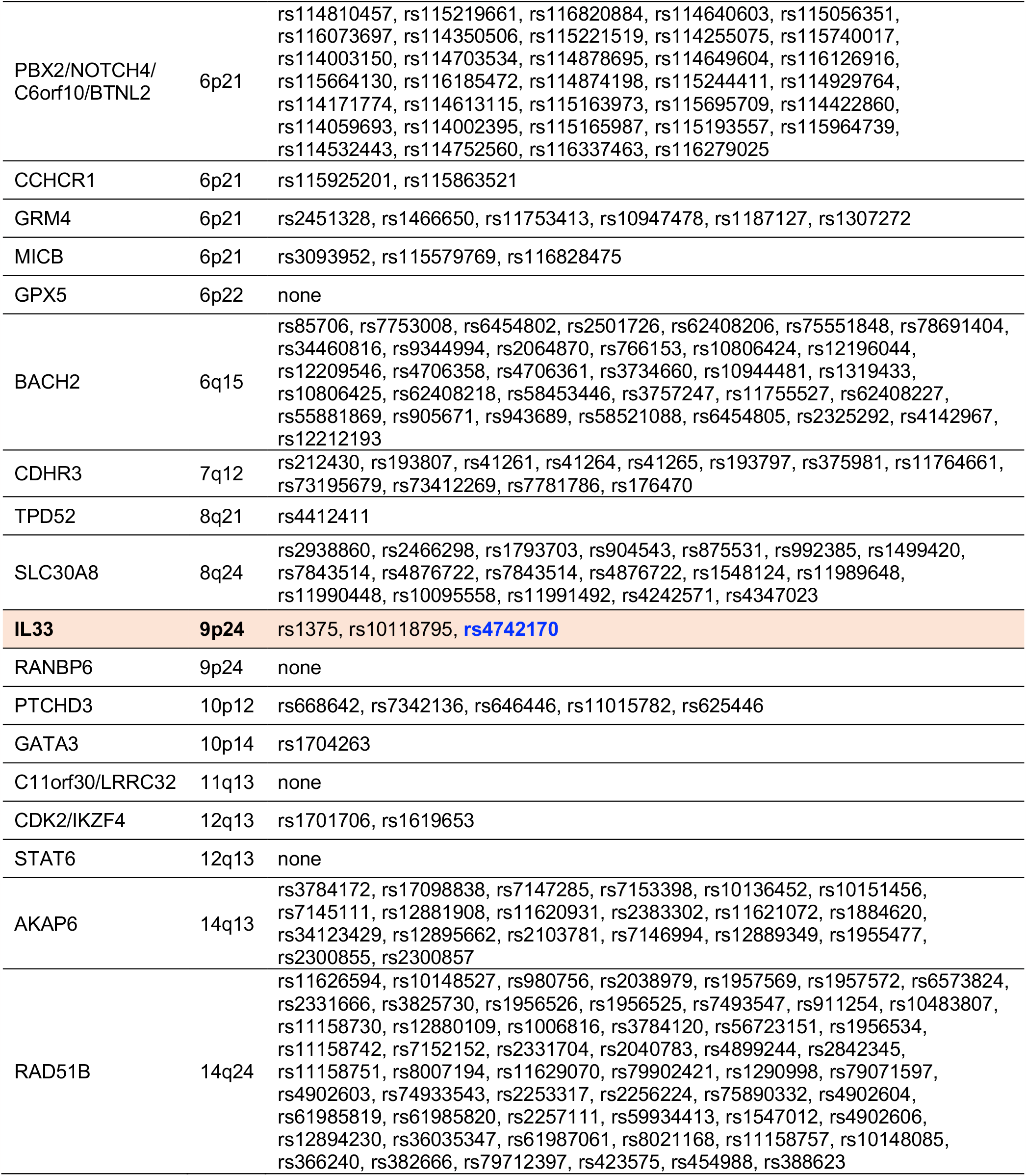
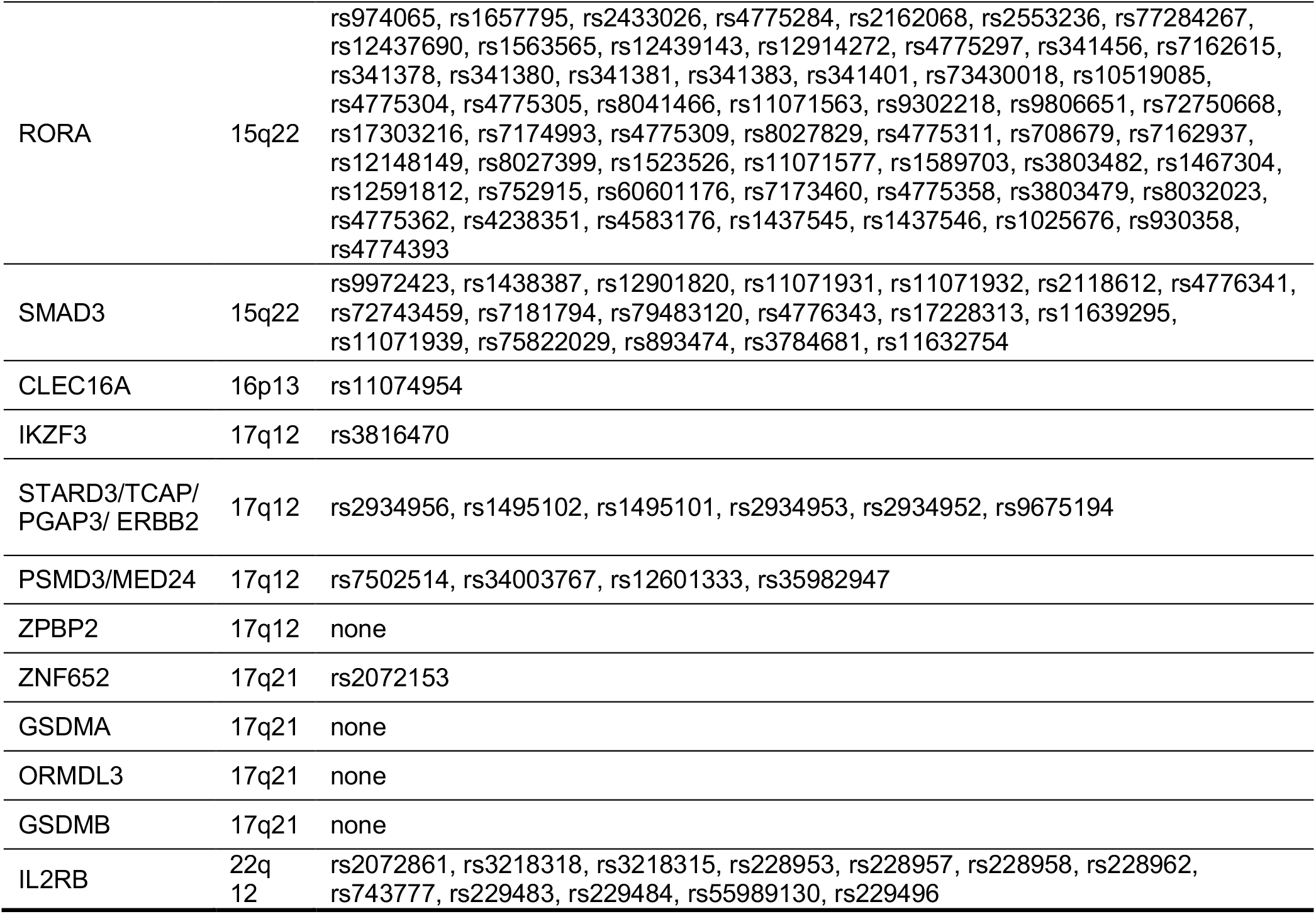
List of asthma GWAS loci assayed, and SNPs identified in the Vindija Neanderthal genome and the French genomes, but not the Yoruba African genome.

**Figure 1.**
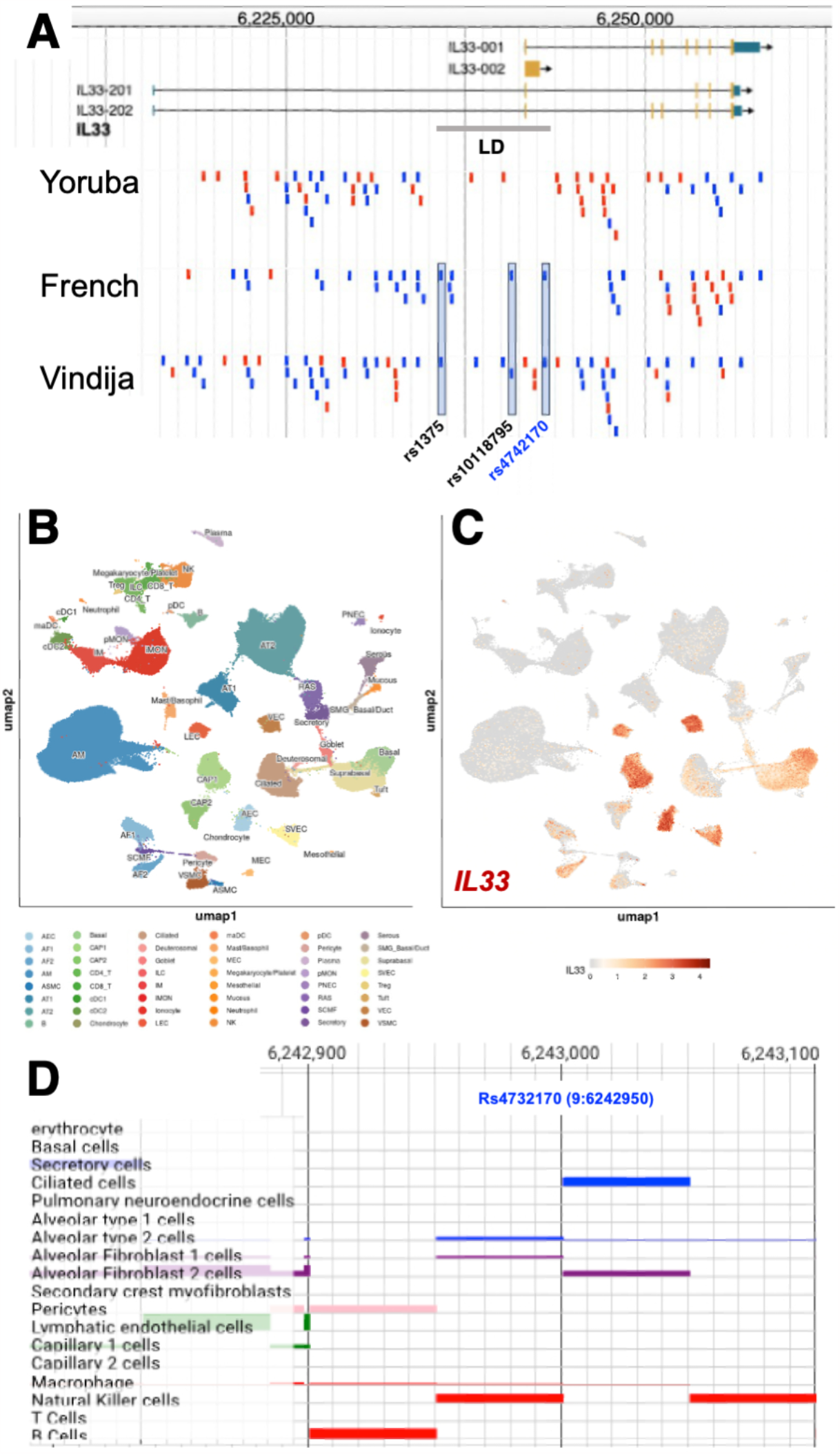
Shared asthma variants in the Neanderthal genome. (A) An example genome browser view showing 3 asthma SNPs (boxed) in *IL33* that are shared between the Vindijia and French genomes, but not the Yoruba genome. LD, linkage equilibrium. (B) A UMAP showing human lung cell clusters based on LungMAP single cell RNAseq data. (C) A feature plot showing *IL33* expression in selected cell types in lung, including airway cell types such as basal cells and ciliated cells. (D) Zoom-in genome track view centered on rs4732170, showing signal in selected cell types including airway ciliated cells.

### Evidence for shared asthma variants to affect gene expression

Most of the Neanderthal-derived SNPs we identified, including those near the lead variants for the asthma GWAS signals, are in non-coding regions of the gene (Table 1). To address if they could change gene expression and contribute quantitively to asthma risk, we screened the list of Neanderthal-derived variants to determine if any known function had previously been ascribed to these variants. We found functional evidence for a Neanderthal-derived variant, rs4742170, that is associated with the *IL33* gene, and has been implicated in childhood wheezing and asthma (27-29). *IL33* encodes an alarmin that serves as a key upstream positive regulator of asthma responses (30). For rs4742170, we identified that the Neanderthal-originated T allele is the disease risk allele. It was shown in a published study that the rs4742170 T allele disrupts repressive glucocorticoid receptor binding to a putative enhancer, leading to an increase in *IL33* enhancer activity (27). Excess IL-33 levels are sufficient to cause asthma phenotypes (31, 32). These functional studies focused on the rs4742170 contribution to asthma, and we add to their findings the origin of the variant allele in the Neanderthal lineage.

### Evidence for association of shared asthma variants in open chromatin regions in human lung cell types implicated in asthma

To gather further evidence that the variant implicated above could alter *IL33* gene expression, we investigated whether they reside in accessible chromatin regions in human lung cell types that actively transcribe *IL33*. Based on publicly available single cell RNAseq data curated by the LungMAP consortium, *IL33* is expressed in a subset of human lung cell types, including a number of endothelial cell types, and basal as well as ciliated cells in the airway epithelium. While the available single nucleus ATACseq data from LungMAP remain sparse, rs4742170 (9:6242950) appears to overlap a region that is accessible in ciliated cells among other cell types. Since asthma is an airway disease, we hypothesize that similar Neanderthal-originated variants could alter transcription factor binding in regions of open chromatin in the airway epithelium to affect expression of genes such as *IL33* and risk of asthma.

## DISCUSSION

In this study, we searched in 51 asthma loci from GWAS and show that 39 of them harbor variants that are present in the Vindija Neanderthal genome and French genome, but not in the Yoruba genome, providing evidence that these variants originated from Neanderthals. In candidate gene based functional studies, one of the variants has been shown to alter the enhancer activity of *IL33*, a key driver of asthma. The identification of the Neanderthal allele in *IL33* offers proof-of-concept for future studies. It is possible that non-biased systematic functional screens across these Neanderthal-originated SNPs may yield additional variants that contribute to the asthma trait.

The discovery and continued refinement of the collective Neanderthal genomes offer tremendous opportunities to trace the origins of human disease (1, 3, 22). Rich data support bi-directional genomic introgression because of Neanderthal-human interbreeding ∼60,000 years ago (33). Evolutionary pressure applied on those genetic traits results in variants that now remain in modern human genomes. It is interesting that many of the genetic traits that have survived the selection are linked to immunity. Our findings here further substantiate this trend by adding asthma to the list. Type 2 immunity, which is at the center of asthma pathology, can be traced back to host reaction to helminth worm infections (34, 35). Specifically, type 2 immunity plays an essential role in clearing the worm (36). A large amount of evidence demonstrates that the asthma response to allergens such as pollen is a byproduct of such heightened type 2 immunity (34, 35). A likely reason that this modern disease trait was selected for is because it conferred immunity against infection in ancient times.

To substantiate disease origin, it is critical to demonstrate that the shared variants change function and/or expression. For rs4742170, functional evidence suggests that it can alter *IL33* enhancer activity (27). Outside of such a candidate-based approach, a non-biased screen is needed to narrow down the remaining variants. One approach is to focus our search to open chromatin regions in disease-relevant cell types, for example, by analyzing publicly available single nucleus ATAC-seq data from the human lung. The proximity of Neanderthal-derived SNPs to lead asthma GWAS SNPs could also be investigated. However, functional tests are ultimately needed to demonstrate causality. With the development of technologies such as massively parallel reporter assay (37, 38), one can devise a non-biased screen to test the potential function of these Neanderthal-associated asthma variants.

Our findings here not only add asthma to the list of diseases that could be traced back to Neanderthals, but they also contribute to our overall understanding of the origin of human diseases. GWAS studies across asthma, allergic rhinitis and eczema have identified shared genetic risk factors (39). These data, together with our results here, support the broad hypothesis that general allergic disease phenotypes originated in Neanderthals.

## METHODS

### Identification of asthma variants in the Neanderthal genome

We carried out a comparative analysis of the single nucleotide polymorphisms (SNPs) in the 51 non-HLA loci that have been associated with asthma by GWAS (17). Using the archaic genome browser (https://bioinf.eva.mpg.de/jbrowse/), we analyzed SNPs from three specific populations, the Vindija Neanderthal ancient genomes, and the French (HGDP00533) and the Yoruba (HGDP00936) from the 1000 genomes project. These three populations were selected to facilitate our search for Neanderthal-derived alleles. This signature is characterized by a variant that is present in both the Vindija and French genomes but absent from the Yoruba genome as shown previously (2, 4, 11).

### Compare variant position to accessible chromatin regions

To identify SNP’s that could impact gene transcription, we analyzed publicly available single cell RNAseq and single nucleus ATACseq data of the human lung on LungMAP.net. We first queried gene expression in single cell RNAseq to assess the cell types that express the gene. We then queried if a SNP resides in an open chromatin region near the targeted gene in the cell types where the gene is expressed.

